# Equal performance but distinct behaviors: *Astatotilapia burtoni* sex differences in a novel object recognition task and spatial maze

**DOI:** 10.1101/2020.08.03.234658

**Authors:** Kelly J. Wallace, Hans A. Hofmann

## Abstract

Sex differences in behavior and cognition can be driven by differential selection pressures from the environment and in the underlying neuromolecular mechanisms of decision-making. The highly social cichlid fish *Astatotilapia burtoni* exhibits dynamic and complex social hierarchies, yet explicit cognitive testing (outside of social contexts) and investigations of sex differences in cognition have yet to be fully explored. Here we assessed male and female *A. burtoni* in two cognitive tasks: a novel object recognition task and a spatial task. We hypothesized that males outperform females in a spatial learning task and exhibit more neophilic/exploratory behavior in across both tasks. In the present study we find that both sexes prefer the familiar object in a novel object recognition task, but the time at which they exhibit this preference differs between the sexes. Females more frequently learned the spatial task, exhibiting longer decision latencies and quicker error correction, suggesting a potential speed-accuracy tradeoff.

Furthermore, the sexes differ in space use in both tasks and in a principal component analysis of the spatial task. A model selection analysis finds that preference, approach, and interaction duration in the novel object recognition task that reach a threshold of importance averaged across all models. This work highlights the need to explicitly test for sex differences in cognition to better understand how individuals navigate dynamic social environments.

## Introduction

For humans and non-human animals alike, navigating dynamic social environments is a constant challenge (O’Connell & Hofmann 2011, Taborsky & Oliveira 2012). Individuals must attend to changing signals and cues, remember stimuli and their valence, assess risk, make inferences from often incomplete or ambiguous information, use and update prior associations, and display socially appropriate behaviors in varied contexts including mate choice (Phelps & Ophir 2009, White & Galef 1999, Hebets & Sullivan-Beckers 2010), territorial/dominance aggression (Reichert & Quinn 2017, Bshary & Brown 2014), social foraging (Rapaport & Brown 2008, Thornton & McAuliffe 2006), and nest building (Keagy et al. 2011).

Males and females frequently make different decisions even within the same social environment. Sex differences in cognition, “the mechanisms by which animals acquire, process, store, and act on information from the environment” (Shettleworth 2010) have been described across taxa (rodents: Orsini & Setlow 2017, Dalla & Shors 2009, Rice et al. 2017, songbirds: Titulaer et al. 2012, lizards: Carazo et al. 2014, poeciliid fish: Lucon-Xiccato & Bisazza 2017a,b, Wallace et al. 2020), although males and females do not always differ in this regard (Lucon-Xiccato & Bisazza 2014 and 2016; Healy 1999, Guillette et al. 2009, Etheredge et al. 2018). When present, sex differences in cognition often arise from varying ecological demands and fitness benefits of cognitive abilities such as sexually divergent pressures related to dispersal, predation, and mate choice (Watson & Platt 2008). For example, males and females can differ in foraging patterns (mice: Maille & Schradin 2016, cormorants: Ishikawa & Watanuki 2002, sandperch Sano 1993) and response to predation (poeciliid fish: Magurran & Seghers 1994, Magurran & Nowak 1991). Cognition can additionally impact fitness via reproductive performance directly, such as sexual display quality (e.g. aspects of song quality in songbirds), (Smith et al 2015, Boogert et al. 2011, Farrell et al. 2011) mate preference, (Shohet & Watt 2009, Branch et al. 2019) or responses to sexual conflict (Cummings 2018, Buechel et al. 2016). Furthermore, behavior and decision-making can vary along many axes including time (seasonality: Clayton & Krebs 1995, Yaskin 2011), social status (Stowe et al. 2007, Layton & Fulton 2014), satiety (Caraco 1981), reproductive status (Lynch et al 2005), and social context (Rosati & Hare 2012).

Burton’s mouthbrooder cichlid, *Astatotilapia burtoni,* is a highly social east African cichlid fish that provides a unique system in which to explore sex differences in cognition in a temporally dynamic social system (Hofmann & Fernald 2001, Maruska & Fernald 2018). In this species, both sexes have been shown to navigate socio-ecological demands that fluctuate across time: males bidirectionally shift between a dominant phenotype (colorful, aggressive, territorial) and a subordinate phenotype (drab coloration, submissive, non-territorial) (Fernald & Hirata 1977, Burmeister et al 2005, Maruska & Fernald 2010, Huffman et al 2012), and females can also form complex social hierarchies and display remarkable behavioral plasticity across reproductive states (Fernald & Hirata 1977, Kidd et al. 2013a, White et al. 1996, Renn et al. 2012, Kidd et al. 2013 a,b). Successfully navigating these complex and ever-changing social surroundings requires cognitive abilities such as transitive inference and cognitive flexibility, (Fernald 2014, Alcazar et al. 2014, Grosenick et al. 2007, Desjardins. et al 2012, Weitekamp et al. 2017, Rodriguez-Santiago et al 2020). Furthermore, this species has emerged as a model system for understanding the overlapping neurobiological mechanisms of social behavior and decision-making (Hofmann 2003, O’Connell & Hofmann 2012, Maruska & Fernald 2018). While cognition has been examined in naturalistic communities of *A. burtoni* (see Wood et al. 2011, Rodriguez-Santiago et al 2020), studies that assess cognitive performance independent of social context and across the sexes are lacking. In the present study we compare the sexes in two cognitive tasks to examine whether and how the social environment might influence cognition *outside* of the fast-paced social lives of this species. Assessing cognition outside of social context is critical for understanding how socioecological pressures influence decision-making and behavior more generally independent of ongoing social interactions.

In the present study, we investigated whether *A. burtoni* males and females differed in cognition by assessing individuals in a novel object recognition task (NOR) and a spatial task (SPA). Both tasks require learning - a change in cognitive state or behavior resulting from prior experience (Shettleworth 2010, Staddon 1983, Domjan 2015)- and test abilities that are relevant to *A. burtoni*’s natural history, as males typically establish territories under specific objects (such as plant debris or rocks), to which they reliably return after extensive excursions in search of females found in shoals on the periphery of the lek (Fernald & Hirata 1977). We hypothesized that if we were to find sex differences in cognition in this species they would manifest as greater male performance in the spatial task. We employed here a spatial task, where individuals were tasked with learning a simple route (behind a left barrier or right barrier to reach a social reward). This paradigm is appropriate given that in the natural environment both sexes are often found in shoals, while dominant males remember the location of their territories. Sex differences in spatial learning have been previously described in fish, with most studies reporting males outperforming females (Carazo et al. 2014, Lucon-Xiccato & Bisazza 2017a,b, Fabre et al. 2014, Saucier et al. 2008, Jonasson 2005, but see Healy et al. 1999 which finds no sex difference and Jones et al. 2003 which emphasizes that home range may be a better predictor of spatial performance than sex). Importantly, behavior exhibited during a task can vary by sex independently of cognitive performance (Burns & Rodd 2008, Mamuneas et al. 2015; Lucon-Xiccato & Bisazza 2016; Etheredge et al. 2018; Titulaer 2012, Dougherty & Guillette 2018), for instance due to where males are typically identified as the “bolder” or “more exploratory” sex, often explained by sex-specific variation in predation pressures and dispersal behavior as well as life history tradeoffs (Lucon-Xiccato & Dadda 2016, Han et al. 2015, Harris et al. 2010, Overli et al. 2006, Gagnon et al. 2016, but see Hughes 1968 & Swanson 1966). In concordance with this literature, we hypothesized that across both tasks males would display more neophilic and exploratory behaviors than females. Lastly, we assessed whether behavior and cognition in the two tasks was correlated to the other (Lucon-Xiccato & Bisazza 2017b) via multidimensional analyses. We hypothesized that, since both memory and spatial learning are needed to successfully navigate the *A. burtoni* social environment, both sexes would successfully learn both tasks and additionally would exhibit correlated individual performance across tasks.

## Materials and Methods

### (a) Housing and Initial Community Formation

128 adult *A. burtoni* (64M/64F) were used from a lab population descended from wild-caught stock. Prior to testing, the fish were maintained in naturalistic communities (114-liter aquaria), which allowed for individual variation in dominance and reproductive status. Both prior to and during testing, fish were kept in reverse osmosis water treated with Seachem^®^ Lake Tanganyika salt and bicarbonate buffer. Lights were on a 12:12 light-dark cycle and fish were fed once daily with Omega One Natural Protein Formula Cichlid Flakes. All individuals were tagged with colored plastic beads for individual identification. One week prior to testing, fish were placed as a community of 8M/8F (n= 8 replicate communities) in a circular pool (1m diameter, 25cm depth, lined with a PEVA curtain liner and Spectrastone White aquarium gravel) containing four halved terra-cotta pots to be used as territorial shelters (Supplemental Figure 1), and at this time fish were weighed and measured for standard length (average standard length: males = 4.78cm, females = 4.76cm). One week after completion of the experiment, individuals were again weighed and subsequently dissected for gonadal weight to calculate gonadosomatic index (GSI, i.e., the ratio of gonad mass to body mass), a proxy for reproductive status.

### (b) Novel Object Recognition Task (NOR)

To assess memory, we utilized a novel object recognition task (also referred to as the spontaneous object recognition task). The novel object recognition task, at its simplest, requires an individual to discriminate between a previously encountered object and a novel one. This paradigm is commonly used in the rodent literature to assess memory (Ennaceur & Delacour 1988, Broadbent et al. 2010), but additionally has the capacity to characterize recognition, neophobia, and exploration (Antunes & Biala 2012).

In the novel object recognition task conducted here, individuals were assessed for relative preference (physical proximity) to a familiarized object versus a novel object in a 114 liter acrylic aquarium filled to a depth of 20cm with a central alley way and two object regions (Figure 1A, B). Four objects were used as stimuli in the task: a pink centrifuge tube, a purple falcon tube cap, a blue weight boat (quartered), and a yellow table tennis ball filled with sand (Supplemental Figure 2). Object pairs and their roles (familiar vs novel) were counter-balanced across communities. Prior to testing, the entire community was habituated at once to two identical objects (coined the “familiar object”) for one hour (e.g. A + A). We chose this method of habituation for two reasons: First, to maintain daily consistency in the timing of the experimental design; and second, to avoid disruption of the social hierarchy, which frequently occurs when only one individual is removed from the community at a time. Approximately 24 hours later, each fish was individually presented with a familiar object and a novel object at the opposite ends of the aquarium (the side of the familiar object, left or right, was counter-balanced across communities) (e.g. A + B). Previous studies in zebrafish have shown that the delay between habituation and testing can range between 2 and 24 hours without impacts on resulting performance (Lucon-Xiccato & Dadda 2014). During the trial, individuals were acclimated in an opaque PVC tube for one minute then allowed to swim freely in the apparatus for twelve minutes. Following the trial, individuals were kept in a holding area until all community members were tested, then simultaneously returned to the community to prevent social hierarchy destabilization. We recorded fish as they transited between the following zones (Figure 1): the central alley, the interaction area (where the object was located), and the corners associated with each object. For each zone we calculated the proportion of time spent in the zone, the number of entries into the zone, and the time of first entry. Performance was calculated as the relative preference for the familiar object (“NOR familiar preference”): the relative proportion of time spent at the familiar object to time spent at the novel object, where a score of 1 means the entirety of time spent at an object was at the familiar object, a score of 0.5 means equal preference, and of 0 means the entirety of time spent at an object was at the novel object.

**Figure 1.**
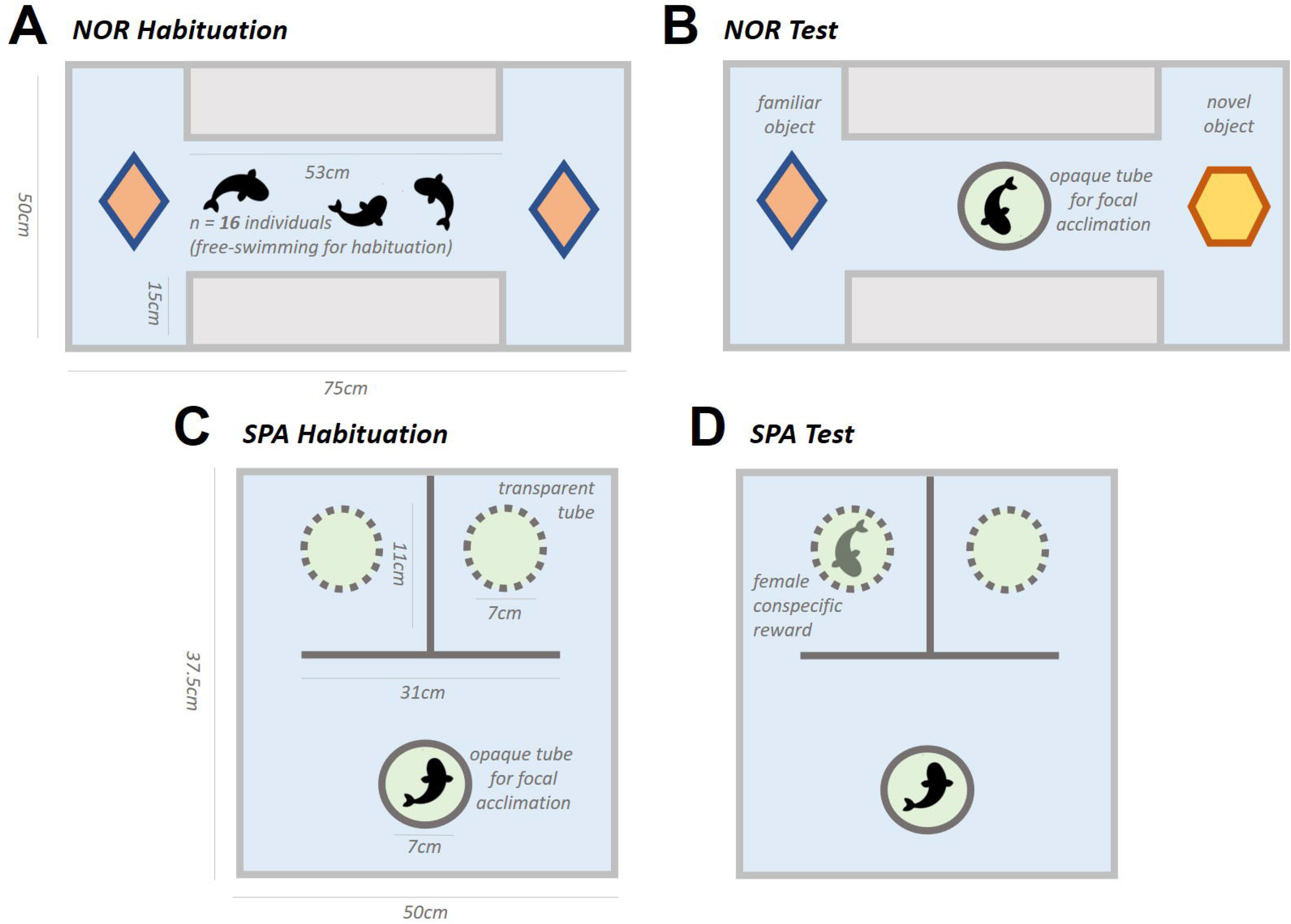
Experimental Designs. The novel object recognition task (A,B) consisted of a 1-hour habituation (community) (A) then a 12-minute test (individual) ~ 24 hours later (B). The spatial task (C,D) consisted of a 1 trial unrewarded habituation (C) then 4 rewarded trials over two days (D).

### (c) Spatial Task (SPA)

Spatial learning involves following a route composed of a sequence of stimuli that must occur in a specific order (Dodson 1988). There has been a long and extensive usage of spatial tasks (“mazes”) to assess spatial learning in fish: see Salas et al. 1996 (goldfish), Lucon-Xiccato & Bisazza 2017a (guppy), Burns & Rodd 2008 (guppy), Jun et al. 2016 (electric fish) Wood et al. 2011 (cichlid). In the spatial task conducted here, individuals in a 57 liter acrylic aquarium filled to a depth of 20cm were required to detour around an opaque barrier in one of two directions (left or right) to reach a social reward consisting of a female in a clear plexiglass tube behind one side of the barrier (Figure 1C, D, average standard length of female conspecific reward = 4.49cm). In *A. burtoni,* both sexes are highly social, and the use of a female reward (from a separate stimulus community and novel to all subjects) for both sexes was employed to avoid potentially confounding effects of male-male aggression. Behind the other side of the barrier, an empty plexiglass tube was placed as a control stimulus. Transit between the rewarded and unrewarded area required an individual to navigate back out in front of the barrier, thus creating a penalty for errors. For all trials, individuals were initially placed in an opaque PVC tube for one minute to acclimate to the aquarium, then allowed to swim freely for twelve minutes. Each individual underwent an initial apparatus habituation trial (using two control stimuli) the day prior to the reward trials. Following this habituation trial, individuals were assayed over four rewarded trials over two days, with the reward stimulus present behind either the left or right barrier, counter-balanced across individuals. Across these trials the same reward female stimuli were used per individual (excluding rare experimental limitations). As with the NOR task, following the trial, individuals were kept in a holding area until all community members were tested then simultaneously returned to the community. We recorded fish as they transited between the following zones (Figure 1): the central neutral area, within 2cm of the various walls, within 2cm of the barrier, the rewarded and unrewarded corners, and the rewarded and unrewarded platforms (which contained the plexiglass tube). For each zone we calculated the proportion of time spent in the zone, the number of entries into the zone, and the time of first entry. To assess performance, we established a learning criterion in which successful learners first approached the rewarded stimulus in all rewarded trials after the first trial (i.e. trials 2-4). Relative preference for the rewarded stimuli (“SPA reward preference”) was calculated as the relative proportion of time spent at the rewarded stimulus to time spent at the unrewarded stimulus, where a score of 1 means the entirety of time spent near a stimulus was at the rewarded stimulus, a score of 0.5 means equal preference, and of 0 means the entirety of time spent at an object was at the unrewarded stimulus.

### (d) Video Scoring

Videos were recorded overhead of the experimental apparat via the Alibi Security Camera system and scored by three human observers (who had not conducted the behavioral assays) in VLC media player using CowLog video scoring software version 3.0.2 (Hänninen & Pastell 2009), a software which has been validated to be highly consistent across multiple observers (Wallace et al. 2020). Focal fish location was recorded though the duration of the test trial of the novel object recognition task and the fourth (test) trial of the spatial task. Additionally, for trials one through three of the spatial task the first approach (rewarded or unrewarded) was recorded.

### (e) Statistics

Data analysis and visualization were performed in R (R Core Team 2013) version 3.6.1 (2019). We conducted an unpaired t-test or unpaired two-sided Wilcoxon Rank Sum test with continuity correction for continuous data split into 2 categories, or a One-way ANOVA or Kruskal-Wallis test for continuous data split into >2 categories (determined by a Shapiro-Wilk normality test). For entirely categorical data we conducted Chi-squared tests. We conducted standard multiple linear regressions to determine correlations and effect sizes of relationships between continuous independent variables using the formula lm(x1 ~ x2) using the R function “lm()”. To identify interaction effects of sex, we conducted linear models with interaction terms using the formula lm(x1 ~ x2 * x3). Effect sizes were calculated as either Cohen’s D using the R function “cohen.d()”in the R package “effsize” (Torchiano 2020) for categorical data or as the Pearson correlation coefficient using the R function “cor()” for continuous data. We conducted principal component analyses (PCA) on z-scored data via the R function “prcomp”. Hierarchical clustering analyses were conducted using the R package “pvclust” (Suzuki et al 2019) with distance metric: “correlation”, linkage metric: “average”, and bootstrap values set to 100. Logistic regression models were constructed and selected using the R package “glmulti” (Calcagno 2020). Variables included in data analysis (including in the principal component analyses) were first assessed via a covariance matrix: any two variables with a Pearson’s correlation coefficient above 0.8 were considered redundant, thus for subsequent analysis the authors used the variable more commonly used in previous literature. Raw data and analysis code can be found at https://github.com/kellyjwallace/Wallace_Hofmann_2020_sex_differences, or via the Texas Data Repository (https://data.tdl.org/).

## Results

### (a) Novel Object Recognition Task (NOR)

Before quantifying performance (first object choice and relative preference for the familiar object) on the NOR task, we first assessed if any objects were preferred independent of familiarity. No object was significantly preferred or avoided (average preferences: cap 55%, weigh boat 45%, ball 43%, tube 57%, Kruskal-Wallis test of preference by object: p = 0.080, df = 3, η^2^ = 0.041, Wilcoxon test of preference between the two pairs: W_41,41_= 389, p = 0.229 and W_55,55_= 497, p = 0.054, respectively). However, because the ANOVA across object and the ball/tube pair showed a near significant bias, we additionally tested whether individuals’ first choice (familiar or novel) differed by the pair of objects they received and found no significant difference (chi^2^ test: p = 0.108, n = 97, X^2^ = 2.577, df = 1).

We then asked whether males and females differed in task performance. We quantified the proportion of individuals that first approached the familiar object in each sex and found that neither sex first approached the familiar object more than would be expected by chance: 63 individuals (38M/25F) first approached the familiar object, and 47 individuals (20M/27F) first approached the novel object. Consistent with this result, we also found that a continuous measure of performance across the length of the trial, the “NOR familiar preference,” did not differ by sex (average NOR familiar preference across both sexes = 0.52, Wilcoxon test: p = 0.386, W_64,64_ = 1268, Cohen’s D = 0.188) and did not diverge from a random 50% expectation in either sex. This preference score did not differ by round (rounds differed in the object pair presented) (Kruskal-Wallis test: p = 0.174, df = 7, η^2^ = 0.038), and was not correlated with GSI in either sex (female linear regression: R^2^ = −0.021, F_37_ = 6.321, p = 0.647, male linear regression: R^2^ = −0.020, F_21_ = 0.565, p = 0.461).

Given that neither sex displayed a significant preference when averaged across the entire task (contrary to our hypothesis that both sexes would learn the task), we further asked whether a preference for either the familiar or novel object could be observed at any time during the task. We initially observed that males exhibited a significant preference for the familiar object in the first minute of the task (Wilcoxon test: p = 0.040, V = 742, n = 49, Figure 2A), and upon a more in-depth analysis found that preference was significantly correlated with time for both males (linear regression: R^2^ = 0.372, F_8_ = 6.321, p = 0.036) and females (linear regression: R^2^ = 0.570, F_8_ = 12.93, p = 0.007), with a significant interaction effect of sex: male’s familiar object preference decreased across time whereas female’s increased (linear regression: R^2^ = 0.575, F_16_ = 9.55, p = 0.0009) (Figure 2B).

**Figure 2.**
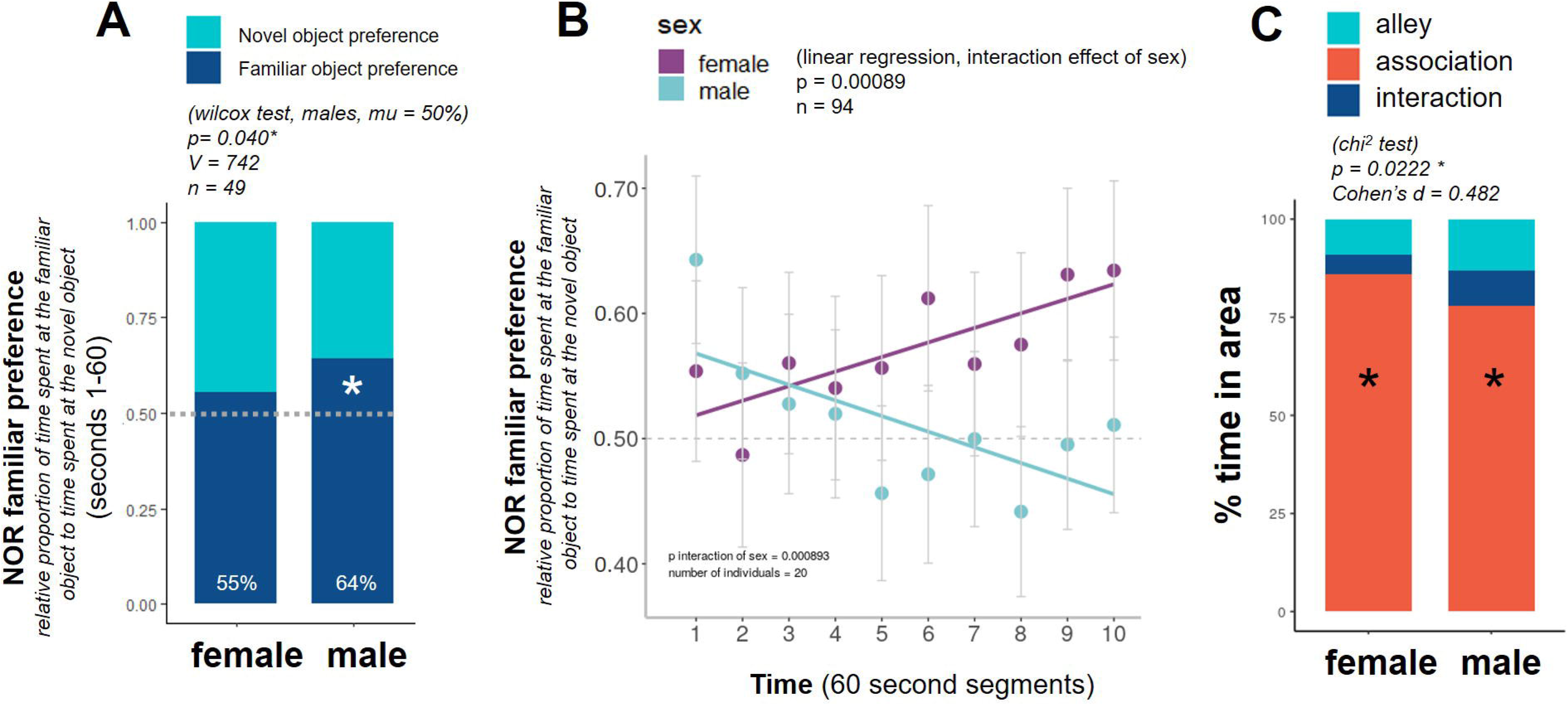
Sex Differences in the Novel Object Recognition Task. Males, but not females, exhibited a significant preference for the familiar object during the first minute of the task (A), and when assessing preference across time male preference decreased over the length of the task whereas female preference increased, with a significant interaction effect of sex (B). Females spent proportionally more time in the object association zone (C).

To assess whether males were the more neophilic sex, we compared latencies to approach the objects and the proportion of time spent with objects: latency to approach either object did not differ by sex (Wilcoxon test: p = 0.463, W_64,64_ = 1275, Cohen’s D = 0.198), nor did the proportion of time spent interacting with either object (Wilcoxon test: p = 0.180, W_64,64_= 987, Cohen’s D = 0.331) though a large proportion of time was spent in the corners near the objects (“association”), significantly more so in females with a small effect (average of females = 85%, average of males = 78%, Wilcoxon test: p = 0.022, W_64,64_ = 1490, Cohen’s D = 0.482) (Figure 2C). Activity (total entries) during the first minute of the task did not significantly differ between the sexes (Wilcoxon test: p = 0.539, W_49,45_ = 1021.5, Cohen’s D = 0.051). Visualizations of focal location across time in the NOR task are available in (Supplemental Figure 3).

We then conducted a multivariate analysis of variables measured in the NOR task (PCA) to further assess relationships between performance and other behaviors observed during the task. The analysis identified 9 PC axes, three of which explained >10% each (and together 74%) of the total variance in the dataset (Supplemental Figure 6, Supplemental Table 1). None of these PC’s differed significantly between the sexes. To identify clusters of behaviors that shared similar loading patterns, we performed a hierarchical clustering analysis of behavior using the PCA eigenvalues. The analysis yielded one highly supported cluster (AU > = 95%): NOR non-engagement (a neophobia metric) and NOR familiar preference.

### (b) Spatial Task (SPA)

To first confirm that no cues (e.g. olfactory) from the reward fish could be detected by the focal fish, we compared the proportion of individuals who first chose the rewarded side during the first trial to a random expectation, and found that individuals first approached the rewarded and unrewarded sides randomly (chi^2^ test: p = 0.313, n = 111, X^2^ = 1.0182, df = 1). To test our hypothesis that the sexes would not differ in spatial learning or, alternatively, that males would outperform females due to territoriality-related cognitive demands, we assessed performance on the final trial of the task (trial 4) and surprisingly found that 31 of 51 females (61%) first chose the rewarded side, which was significantly more than the 25 of 59 males (42%) who first chose the rewarded side (chi^2^ test: p = 0.043, n = 111, X^2^ = 4.064, df = 1, Supplemental Figure 4). We then assessed performance based on a criterion of learning where we classified learners as those who first approached the rewarded stimulus in all rewarded trials after the first trial (trials 2-4). The probability for this to occur by random choice is 12.5%.

Interestingly, 7 of 58 males (12.1%) reached this criterion (suggesting random choice), whereas 12 of 51 females (23.5%) reach this criterion, representing a significant cumulative binomial probability in which the females learn the task more than would be expected by chance (p = 0.021, Figure 3A). However, this is not a significant sex difference in performance, as the proportion of males that reached the learning criterion did not differ from proportion of females that reached the learning criterion (chi^2^ test: p = 0.186, n = 109, X^2^ = 1.744, df = 1).

**Figure 3.**
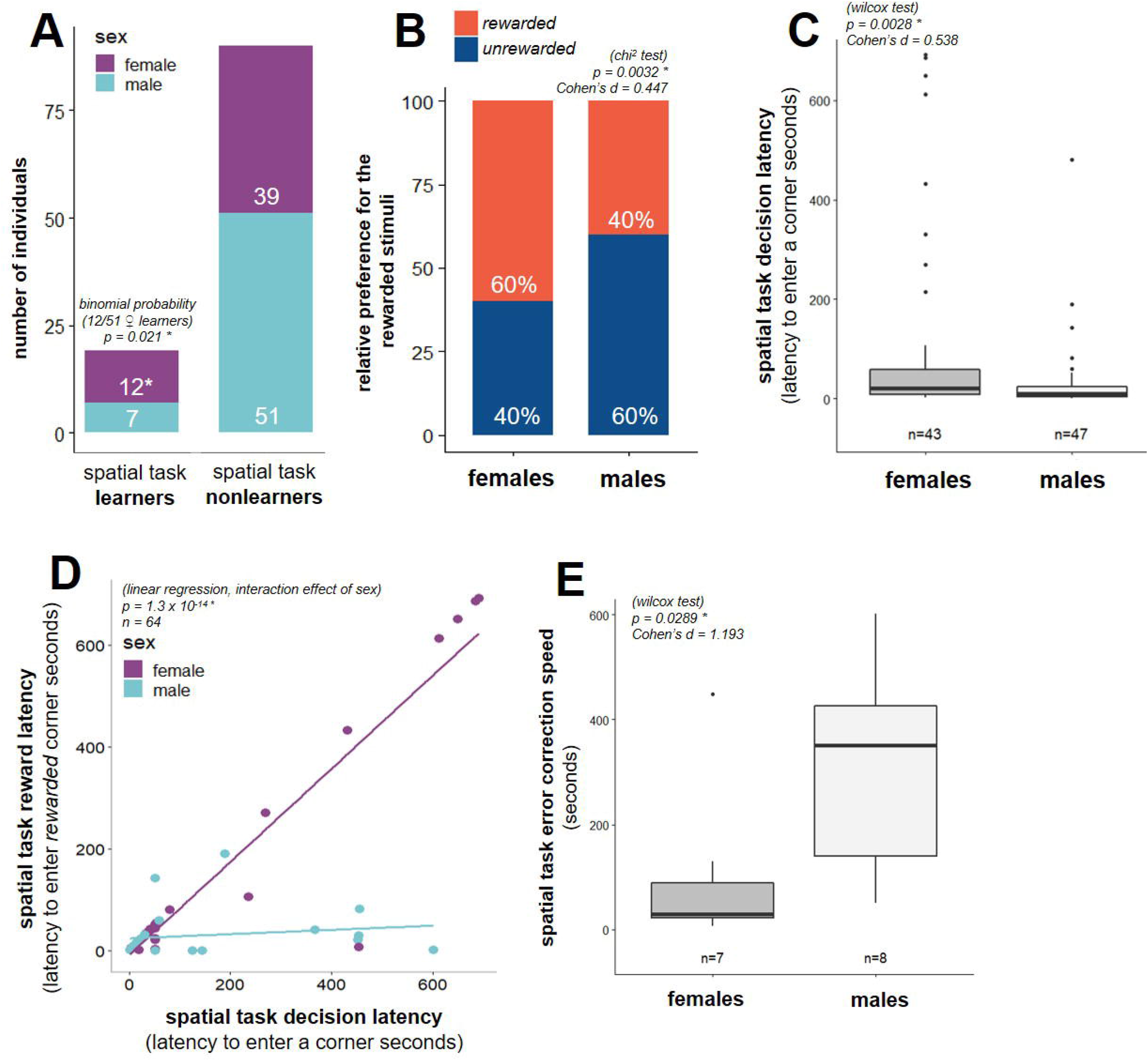
Sex differences in the Spatial Task. More females reached the spatial task learning criterion than would be expected by chance, whereas males did not (A). Females exhibited a greater preference for the reward stimuli during the task (B). Females also demonstrated longer decision latencies (C) and their decision latencies significantly predicted their latency to solve the task (enter the reward area) whereas this relationship was not seen in males (D). When subsetting individuals who first entered the unrewarded side and then corrected and entered the rewarded side, females’ error correction speed was faster than males’ (E).

Next, we asked whether males were the more neophilic sex and whether behavior was related to performance in the task. To this end, we assessed several behavioral and performance metrics in the fourth trial of the assay. Interestingly, we detect a small effect where *males* spent significantly more time spent near the walls of the aquarium during the task (thigmotaxis), which is commonly considered an anxiety behavior (Maximino et al. 2010) (Wilcoxon test: p = 0.045, W_53,50_= 1023, Cohen’s D = 0.435). We found a small effect where females exhibited a significantly higher SPA reward preference across the duration of the task (female average = 0.6, male average = 0.4, Wilcoxon test: p = 0.032, W_64,64_ = 1346, Cohen’s D = 0.447) (Figure 3B). This preference was not correlated with GSI in either sex (female linear regression: R^2^ = −0.006, F_38_ = 0.770, p = 0.386, male linear regression: R^2^ = −0.026, F_24_ = 6.321, p = 0.549). Additionally, females took significantly *more* time to decide than males with a moderate effect size (Wilcoxon test: p = 0.0028, W_64,64_= 1381, Cohen’s D = 0.538) (Figure 3C). Also, their latency to solve the task (enter the rewarded area) was highly correlated with this latency to decide, a relationship that was not present in males (linear regression: R^2^ = 0.851, F_60_ = 121.3, p = 1.296×10-14) (Figure 3D), suggesting that the sexes might differ in error correction. To test this idea, we subsetted the results by individuals who first approached the unrewarded side and subsequently entered the rewarded side and measured the time between the first unrewarded entry and the subsequent rewarded entry. We found that, indeed, females correct these first approach “errors” significantly faster than males with a very large effect (female average = 101 seconds, male average = 309 seconds, Wilcoxon test: p = 0.029, W_7,8_ = 9, n = 15, Cohen’s D = 1.193) (Figure 3E). Latency to solve the task was not correlated with GSI in either sex (female linear regression: R^2^ = 0.057, F_33_ = 3.038, p = 0.091, male linear regression: R^2^ = −0.10, F_21_ = 0.0002, p = 0.990). Visualizations of focal location across time in the SPA task are available in (Supplemental Figure 5).

As with the NOR task, we conducted a principal component analysis of variables measured in the spatial task to understand the relationships between the behaviors observed during the task. Of the 10 PC axes identified, we explored further the four PCs that explained >10% each (and together 85%) of the total variance in the dataset (Supplemental Figure 7A, Supplemental Table 2). Interaction, association, decision latency, and interaction latency loaded most strongly onto PC2 (20% of the variance) and significantly differed by sex with a large effect (Wilcoxon test with FDR correction, p =0.008, W_26,24_ = 481, Cohen’s D = 0.884) (Supplemental Figure 7B). None of the other PCs varied across sexes, and in a hierarchical clustering analysis of the eigenvalue patterns did not yield any well supported clusters (AU > = 95%).

### (d) Relationships across the tasks

Completing the NOR task requires learning and memory, and both abilities are likely important for successfully navigating the cognitive demands of *A. burtoni* social life. We therefore asked whether performance across the two tasks was correlated. When first simply using the categorical metric of “first choice” (rewarded or unrewarded in SPA, familiar or novel in NOR) the two tasks are not correlated. However, performance in the two tasks can be compared appropriately only for those individuals who entered both the familiar and novel object recognition task areas *and* the rewarded and unrewarded areas of the spatial task. 13 males and 7 females met this criterion. When we then correlated the continuous preference metrics across the tasks (relative preference for the familiar object in the novel object recognition task and relative preference for the rewarded stimuli in the spatial task), we found that they were indeed significantly correlated in a sex-invariant manner (linear regression: R^2^ = 0.229, F_18_ = 5.359, p = 0.032) (Figure 4A). Because we had previously discovered that decision latencies differed by sex in the SPA tasks (Figure 3C), we next asked whether decision latencies related to choices across tasks, and observed a modest effect where individuals who first approached the novel object in NOR task showed longer decision latencies in SPA (Wilcoxon test: p = 0.033, W_58,39_= 427, Cohen’s D = 0.333), and conversely, individuals who first approach the rewarded stimulus in SPA exhibit longer decision latencies in NOR (Wilcoxon test: p = 0.010, W_53,50_ = 1062, Cohen’s D = 0.029) (Figure 4B,C).

**Figure 4.**
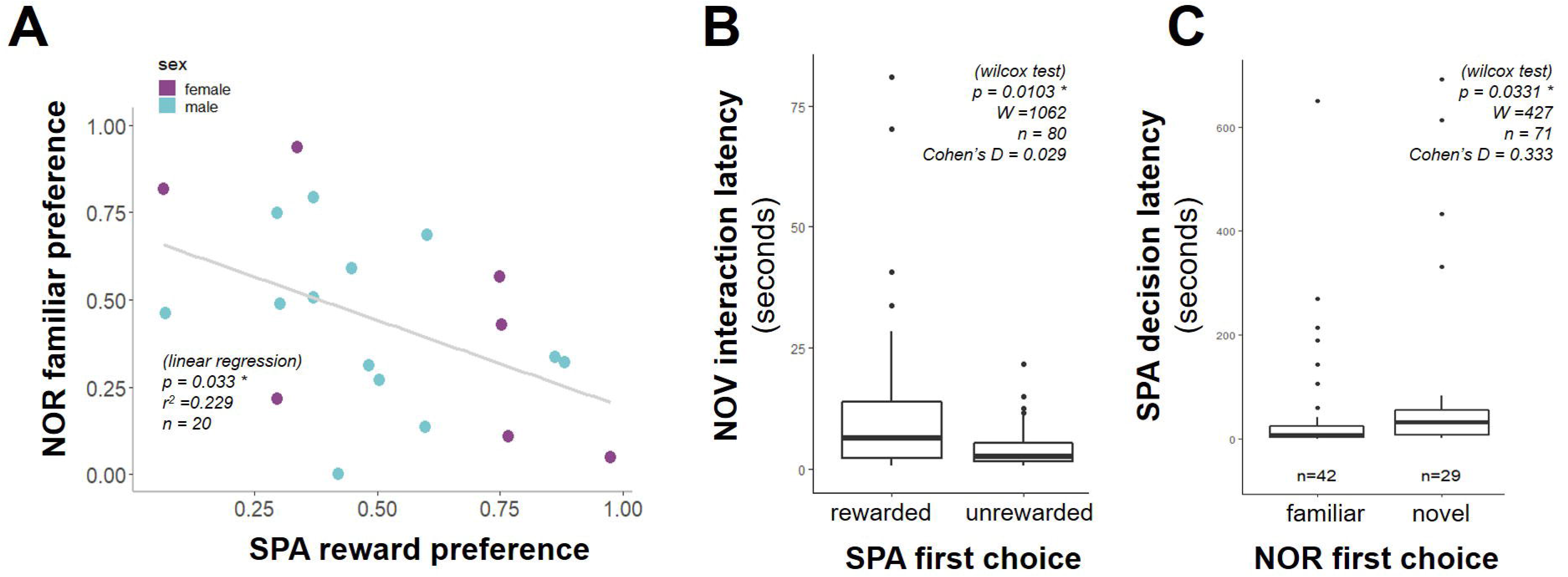
Individual Variables are Related Across Tasks. When subsetting by individuals who had an intermediate level of preference, relative preference scores are correlated across tasks (A). Decision latency in the NOR task differed by the first choice made in the SPA task (rewarded or unrewarded) (B), and decision latency in the SPA task differed by the first choice made in the NOR task (familiar or novel) (C).

To better understand the relationships between variables across both tasks, we conducted a principal component analysis that included the variables from both the SPA and NOR tasks (Figure 5, Supplemental Table 3), and identified four principal components (of ten total) that explained at least 10% each (and together 61%) of the total variance. The first two principal components were largely defined by task: PC1 (accounting for 24% of the variance) loaded most strongly with measures of activity and reward preference in the spatial task, whereas PC2 (accounting for 16% of the variance) loaded primarily by object preference and approaches, though it also included latency variables across both tasks. PC3 (11%) loaded most strongly by variables associated with stimulus interaction across both tasks, and PC4 (10%) loaded primarily by a single variable: SPA reward latency. No PC axes in this combined analysis differed significantly by sex. When clustering behaviors by their eigenvalue patterns in this PCA, we identified five highly supported clusters (AU > = 95%), some of which were with a task and some which spanned across tasks (Figure 5, highlighted in grey boxes).

**Figure 5.**
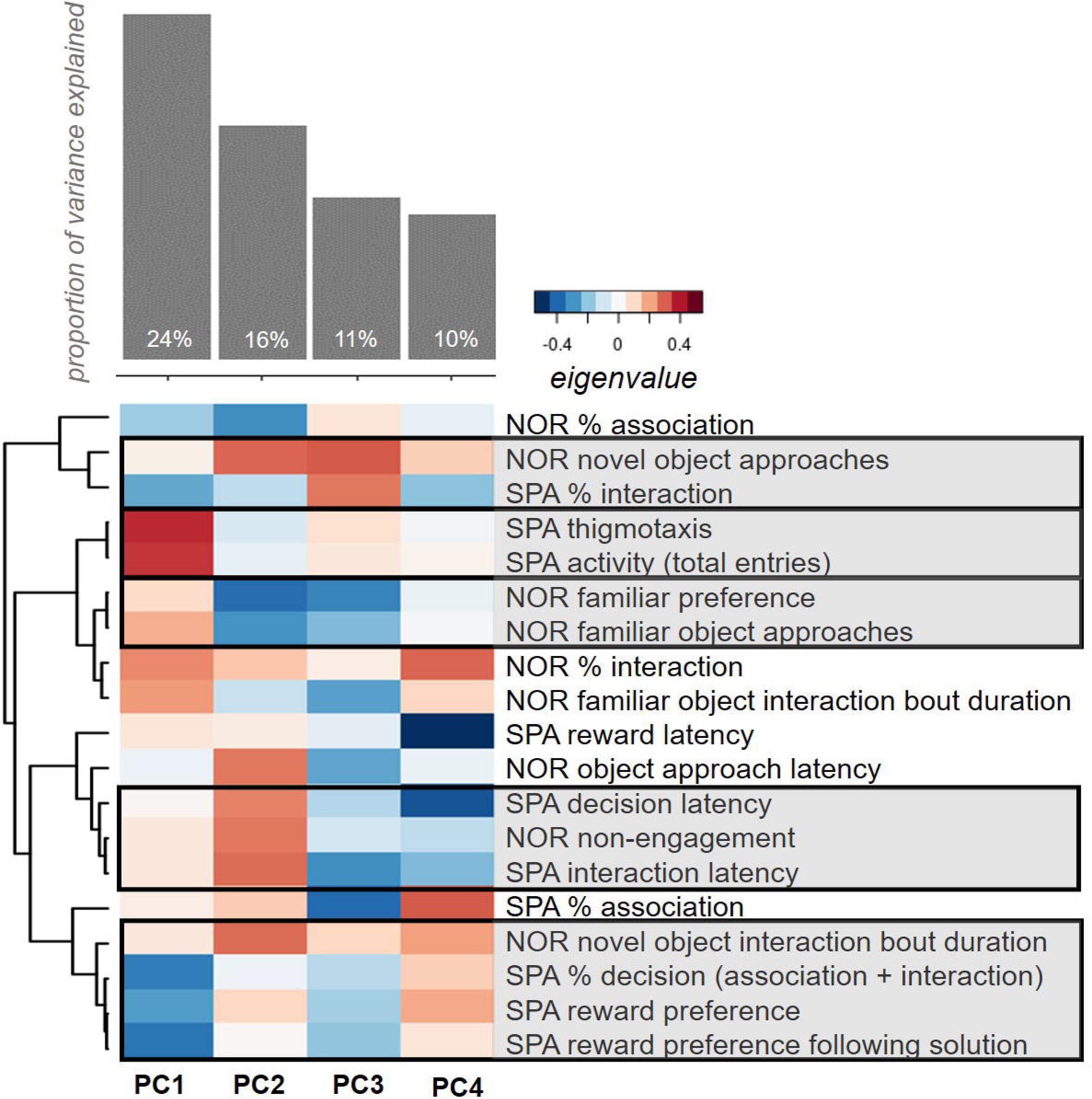
Principal Component Analysis of Variables from Both Tasks. A PCA of variables assessed in both tasks identifies four axes that explain >10% of the variance. Shown here are the proportion of variance for each PC axis (top) and a heatmap of the eigenvalues of each behavioral variable for each PC axis (bottom) (A). The first PC axis (24%) can be described by measurements of activity and preference in the spatial task. The second PC axis (16%) load most strongly by preference in the novel object recognition task as well as measurements of latency across both tasks. The third PC axis (11%) loads most strongly with interaction variables across both tasks. The fourth PC axis (10%) loads very strongly by a single variables: SPA latency to solve the task. A hierarchical clustering of variables by loading patterns (dendrogram) identified five highly supported clusters (AU > = 95%, grey boxes), three of which contain variables form both tasks.

Finally, to identify which variables across both tasks were best able to distinguish males and females, we conducted three logistic regression model selection analyses to predict sex via an exhaustive screening of all possible models and selecting based on AICc: one with variables from the NOR task, one with variables from the SPA task, and a combined model. Using variables from the NOR task, the selected model (of 511 possible models, McFadden’s Pseudo-R^2^ value of 0.032) included only NOR novel object interaction bout duration (Supplemental Table 4). Using variables from only the SPA task, the selected model (of 1023 possible models, McFadden’s Pseudo-R^2^ value of 0.213) included the SPA preference measure and % association (Supplemental Table 5). For each of analysis we additionally conducted a model-averaged importance of terms analysis. In a model-averaged importance of terms analysis, the importance of a term is determined by the number of models the term appears in and their weights (a metric of the model’s likelihood to be the “best” model). Neither of these task-specific models identified a term that reached the importance threshold of 0.8 (Supplemental Tables 4,5). Using variables from both tasks, the selected model (of 524,287 possible models, McFadden’s Pseudo-R^2^ value of 1.00) included the following variables from both tasks: NOR novel object interaction bout duration, NOR familiar preference, NOR novel object approaches, NOR association, NOR interaction, NOR object approach latency, SPA decision latency, SPA % association, SPA reward latency, NOR non-engagement, SPA reward preference, SPA reward preference following solution, and SPA thigmotaxis (Table 1, bolded). The terms in a model-averaged importance of terms analysis that reached the importance threshold were three NOR variables: NOR novel object interaction bout duration (0.994), NOR familiar preference (0.938), and NOR novel object approaches (0.927). (Table 1, highlighted with an asterisk)

**Table 1.** Model Selection and Model-Averaged Importance of Terms Analysis. Through an exhaustive “all-possible models” selection analysis, we identified the model with the lowest AICc value, which included 13 variables spanning both tasks (variables in the resulting model in bold). In a model-averaged importance of terms analysis (in which each term’s importance score is related to the number of models it appears in and the weight of those models), three NOR variables reach the importance threshold (highlighted with asterisks).

| Term Name | Importance Score |
| --- | --- |
| **NOR novel object interaction bout duration** | 0.994 * |
| **NOR familiar preference** | 0.938 * |
| **NOR novel object approaches** | 0.927 * |
| **SPA decision latency** | 0.800 |
| **SPA % association** | 0.769 |
| **NOR % interaction** | 0.758 |
| **SPA reward latency** | 0.622 |
| **NOR non-engagement** | 0.548 |
| **NOR % association** | 0.520 |
| **SPA reward preference** | 0.394 |
| SPA % decision (association + interaction) | 0.365 |
| **SPA reward preference following solution** | 0.358 |
| SPA % interaction | 0.352 |
| NOR familiar object interaction bout duration | 0.281 |
| **SPA thigmotaxis** | 0.239 |
| **NOR object approach latency** | 0.183 |
| SPA interaction latency | 0.161 |
| SPA activity | 0.096 |
| NOR familiar object approaches | 0.033 |

## Discussion

The present study investigated sex differences in cognition (outside of social contexts) in a species in which both sexes navigate temporally dynamic socio-ecological pressures would show less sex differences in cognition. We used Burton’s mouthbrooder cichlid, *Astatotilapia burtoni,* a species well-characterized for its fast-paced and complex social system (Hofmann & Fernald 2001). We assessed male and female *A. burtoni* (across 8 replicate communities, a total of 128 individuals) in two cognitive tasks: a novel object recognition task and a spatial task. In the novel object recognition task, both sexes exhibited a preference for the familiar object but at different times of the task, and males spent less time associating with the objects than females. In the spatial task, females reached a learning criterion above chance levels and exhibited stronger preferences for the reward stimulus. Females exhibited longer decision latencies and quicker error correction speeds. These differences were also captured in a principal component analysis of the spatial task, which identified a PC axis that significantly varied by sex and loaded most strongly by stimulus interaction and latency. Broadening our analysis to look at relationships across both tasks, we found that in a sex-invariant manner preference scores across the two tasks significantly correlated to each other (preference for the familiar object in the novel object recognition task, and preference for the rewarded stimulus in the spatial task), and decision latencies predicted first choice behaviors across tasks. The first axis of a principal component analysis using variables from both tasks was primarily loaded by SPA preference and activity. A hierarchical clustering analysis of eigenvalue patterns validated behavioral relationships both within and across tasks, and additionally identified a potential neophobia relationship across the tasks. Finally, using a logistic regression model selection approach, we found that the behavioral variables that best predicted sex spanned both tasks. However, when assessing the importance of variables across all models, only NOR measures (novel object interaction bout duration, familiar preference, and number of novel object approaches) reached an importance threshold.

### (a) Sex differences in the Novel Object Recognition Task

In the novel object recognition task, both sexes exhibited a significant preference for the familiar object, confirming our hypothesis that both sexes would learn the task. While rodents (of both sexes) typically prefer the *novel* object, studies in fish have yielded conflicting results (May et al. 2016, guppies: Miletto Petrazzini et al. 2012, Lucon-Xiccato & Dadda 2016). Where we did identify sex differences was in the *timing* of the NOR preference for the familiar object: males exhibited the familiar preference early on in the task, whereas females exhibited the familiar preference later in the task. Studies in fish have found cases where preferences are only exhibited during for the first few minutes of the task (Lucon-Xiccato & Dadda 2016, Miletto Petrazzini et al. 2012). We do not believe this sex difference in the timing of the familiar preference in our experiment to be a reflection of motivation or initial anxiety during the task, as latency to approach either object did not differ by sex. Furthermore, both sexes are equally active in the first minute of the task (our closest proxy to freezing behavior, Blaser et al. 2010). Instead, we suggest that this timing difference reflects variation in space use between the sexes. In a NOR experiment conducted by Lucon-Xiccato & Dadda (2016), zebrafish males exhibited similar object preferences to females, but more quickly approached the novel object. The authors emphasize, as we do, that the differences found likely do not reflect differences in memory but rather space use (as zebrafish and *A. burtoni* are both small, social fish that may respond similarly to novel isolated environments).

One additional consideration that may explain the sex difference in NOR familiar preference is that, in our experimental design, the animals were familiarized to the task (all individuals from a given community were familiarized together). A similar study in juvenile guppies also used a NOR task while minimizing social isolation (i.e., habituation to the apparatus occurred in pairs) and found a preference for the familiar object during the subsequent test (Miletto Petrazzini et al. 2012). In ravens, dominance status influenced approach latencies in a novel object task, and this relationship between dominance status and approach varied by social context (e.g. same-sex vs mixed-sex pair) (Stowe et al. 2006). Dominance status has been shown to relate to exploratory behavior (Dingemanse & de Goede 2004, Stapley & Keogh 2004, Carazo et al. 2014) and cognitive performance across multiple tasks and taxa (pheasants, Madden et al. 2018, starlings, Boogert et al. 2006, dogs, Pongracz et al. 2008). In *A. burtoni*, males vary in dominance status, with dominant males aggressively establishing and defending territories (Fernald & Hirata 1977). Because the habituation phase of the NOR task was conducted in a social setting, the preferences exhibited during the test may be the result of social associations (e.g. males more readily seeking the familiar object as a territory). While we failed to discover a relationship between GSI (as a proxy for reproductive state), further experiments assessing within-sex variation in *A. burtoni* and a possible role of dominance and reproductive status are needed to identify potential social factors driving NOR preferences.

### (b) Sex differences in the Spatial Task

Across both sexes only a small proportion of individuals successfully reached the learning criterion in the spatial task. This is not unexpected, given that our experimental paradigm has few trials and a strict learning criterion (“learners” had to learn after only one training trial and not make a mistake in the following three trials). A similar design has been used in another cichlid species, the Nile tilapia *Oreochromis niloticus* (Lombardi Brandão et al. 2015). However, this study also used a T-shaped barrier in which a focal individual had to navigate around by making either a left turn or right turn to reach a reward and after 30 trials (2x/day) with a criterion of 9/10 correct trials in a row 50% of their focal individual successfully learned the task. In contrast, our design lacked landmark cues and used a much shorter training procedure. It is thus not surprising that only a small proportion of individuals reached the learning criterion.

It has been suggested that in territorial species competition for mates drives increased spatial cognitive performance (Heske & Ostfeld 1990), and several studies have indeed shown that the more territorial sex (which is often male) exhibits increased spatial cognition (Gaulin & Fitzgerald 1989, Carazo et al. 2014, Rice et al. 2017, Carazo et al. 2014, Lucon-Xiccato & Bisazza 2017, Fabre et al. 2014, Saucier et al. 2008, Jonasson 2005, but see Jones et al. 2003, Healy et al. 1999, Lucon-Xiccato & Bisazza 2017a). We therefore hypothesized that because some *A. burtoni* males defend territories (and territory defense is the only scenario in which members of this species exhibit clearly delineated home ranges), males would exhibit greater spatial learning performance. Contrary to our expectations, we found that in the spatial task it was *females* who reached the learning criterion more than would be expected by chance and thus successfully learned the task, whereas males did not. However, males approached either stimulus more quickly than females, suggesting that the sex difference in spatial learning performance was not due to a difference in motivation. Despite females reaching the learning criterion more than expected by chance, the sexes did not significantly differ in performance when compared to each other. Female *A burtoni* must also attend to spatial locales for spawning in male territories, and furthermore females are capable of exhibiting male-typical behavior (including territory defense) in the absence of males (Renn et al. 2012), suggesting that despite rarely exhibiting territory defense under normal conditions the cognitive abilities needed for territory defense are present in both sexes (White et al. 1996). Further research in *A. burtoni* females on foraging ecology, predation pressure, risk assessment, and shoaling behavior is necessary to validate these explanations. And even though reproductive status (GSI) was not significantly correlated with performance in either sex in the present study, future research will need to examine within-sex variation in cognition in this species.

In the PCA of the spatial task, PC2 (explaining 20% of the variance) significantly differed by sex. This PC2 loaded most strongly by variables of stimuli interaction, association, and latencies. Additionally, females exhibited longer latencies to decide in the spatial task with medium effect and more frequently reached the learning criterion. Taken together these results suggest a possible speed-accuracy tradeoff, such that females are the slower but more accurate sex in this task. Speed accuracy tradeoffs have been previously identified in the literature (chickadees: Guillette et al. 2011, voles: Mazza et al. 2018, see Chittka et al. 2009 for review). This interpretation is further supported by the finding that females exhibit their NOR familiar preference later in the task, suggesting that females may be the “slower” sex in the NOR as well. Additionally, females corrected errors much faster than males, suggesting higher accuracy as well as increased cognitive flexibility. Cognitive flexibility is the ability to modify behavior in response to variations in consequences or context of the environment (Bond et al. 2007; Easton 2005). While we did not specifically assess cognitive flexibility in this task, our observations of quicker female error correction (a term used to assess cognitive flexibility in other tasks, see Mazza et al. 2018) complements previous findings that female guppies have been shown to outperform males in tests of cognitive flexibility (Miletto Petrazzini et al. 2017, Lucon-Xiccato & Bisazza 2017). Given the large effect size of the sex difference in error correction found in this study, further experiments in this species quantifying sex differences in cognitive flexibility are likely to yield fruitful insights.

### (c) Relationships across tasks

A growing body of literature has demonstrated a high correlation in performance across cognitive tasks (Chiappe & McDonald 2005, Kolata et al. 2008, Banerjee et al. 2009). At this current state of research on sex differences in (animal) cognition, relatively few studies have specifically investigated sex differences across multiple cognitive tasks. Those that do often find sex differences in specific tasks rather than general greater performance by one sex across all tasks (e.g. Dalla & Shors 2009, Lucon-Xiccato & Bisazza 2017a, Wallace et al. 2020, Miletto Petrazzini et al. 2017; Lucon-Xiccato & Bisazza 2014; Brust et al. 2013; Carazo et al. 2014; Laland & Reader 1999). Because both memory and spatial learning are likely needed by *A. burtoni* males and females to successfully navigate the dynamic and fast-paced social environment typical for this species, we predicted performance would be correlated across tasks for both sexes. In a PCA of both tasks, SPA variables largely loaded on PC1 (explaining 24% of variance) and NOR variables load on PC2, suggesting that these tasks are not strongly correlated. However, preference for the familiar object in the NOR task correlates significantly preference for the reward stimulus in the SPA task at an individual level.

Furthermore, when clustering variables by their eigenvalue patterns in this PCA, a majority of clusters contain variables from both tasks. NOR novel object approaches clustered with SPA interaction, both potential measures reflecting boldness/neophilia. SPA thigmotaxis (wall-hugging) clustered with SPA activity (number of entries during the task), which is unsurprising given both these behaviors reflect anxiety-like behaviors. NOR familiar preference clustered with the number of familiar object approaches, consistent with what we would expect. The two SPA latencies (decision and interaction) clustered with NOR non-engagement, reflecting another possible boldness/anxiety/neophobia cluster. Lastly, NOR novel object interaction bout duration, SPA decision (time spent in both the association and interaction areas), and SPA preference (both over and preference following solution) clustered together. It is important to note that these hierarchical clusters of eigenvalues not only help identify potential dimensions where the sexes might differ; more generally, they can validate the interpretation of our behavioral measurements. For example, the clustering of SPA thigmotaxis and activity confirm our expectation that both these behaviors relate to anxiety (Maximino et al 2010). Additionally, these clusters elucidate some behavioral patterns that may not be as expected: the clustering of NOR novel object approaches and SPA interaction suggests that close contact with the conspecific reward in the SPA should perhaps not be considered a measure of “social affiliation,” rather it may additionally relate to boldness.

Finally, we hypothesized that the spatial task would capture greater sexual dimorphism in cognition than the novel object recognition task, and indeed a PCA of variables in the SPA task identified an axis (PC2) that significantly differs by sex, whereas no axis in the NOR PCA differed significantly. Furthermore, the only sex differences with large effect sizes resulted from the SPA task, suggesting that the sexes diverge in this task more than the NOR task. But counter to this hypothesis, only variables in the NOR task (novel object bout duration, familiar preference, and number of novel object approaches) reached an average importance threshold across all models. These three variables are all related to exploration of a novel object, indicating that variation in neophobia may be at the root of the observed sex differences in *A. burtoni*. Additional experiments that directly quantify exploratory behavior, stress response, and neophobia in this species will help disentangle these behavioral factors from cognitive performance and will facilitate improved cognitive assay design in the future.

### (d) Conclusion

In the present study, we asked to which extent the sexes differ in cognition in a species that lives in a highly dynamic social environment. This question reflects a larger theoretical framework on the reduction or exaggeration of sex differences depending on the social environment. This framework has been successfully applied as it relates to mating system. In voles, monogamous species show a lower degree of sexual dimorphism in morphology, behavior, and cognition relative to their non-monogamous congeners (Kleiman 1977, Jacobs 1995, Heske & Ostfeld 1990, Jacobs et al. 1990, Gaulin & Fitzgerald 1989). And in two species of polygynandrous fish (the shanny and the Azorean rock-pool blenny), females have larger home ranges and a larger dorsolateral telencephalon (hippocampus) than males, though this has yet to be correlated to their learning and memory abilities (Costa et al. 2011). In the present study we hypothesized that in more dynamic or fluctuating social environments we would find fewer sex differences in cognition performance, and indeed found no significant differences in *performance* in either task (though the greater female performance on the spatial task warrants additional work). We did, however, identify several behavioral differences between the sexes primarily related to neophobia, space use, and action latencies. Without a rigorous, phylogenetically controlled species comparison and without a more detailed understanding of the variation in ecologically relevant behaviors (e.g. shoaling, foraging, anti-predation behavior) between the sexes in *A. burtoni,* we cannot definitively know whether the sex differences we have observed here are greater or less than “expected”, but we contribute to this question by uncovering previously undescribed differences in cognition and behavior across the sexes. Our findings were facilitated by an experimental design employing two distinct cognitive tasks followed by multivariate analyses to describe the complex relationships observed in the data. We argue that pairing a more “reductionist” design of discrete suites of cognitive tasks with a strong foundation of more “naturalistic” observations of behavior is a powerful approach to understanding the relationship between the social environment and cognition. Future research across suites of cognitive tasks specifically designed to test variation within each sex and between other social phenotypes (e.g. mating systems, reproductive status, dominance status) will allow us to further relate cognitive performance to ecologically-relevant behaviors. Ultimately, these research endeavors will allow us to better understand the evolution of cognition and decision-making in a dynamic social world.

## Supporting information

Electronic Supplementary Materials

## Data Accessibility

Raw data (Microsoft excel) as well as analysis and visualization R code can be found at https://github.com/kellyjwallace/Wallace_Hofmann_2020_sex_differences, via the Texas Data Repository (https://data.tdl.org/), or via request to the corresponding author.

## Acknowledgements

The authors would like to thank Kavyaa Choudhary, Layla Kutty, Don Le, Matthew Lee, and Karleen Wu for assistance in data collection, and to Mike Ryan, Tessa Solomon-Lane, Felicity Muth, and all members of the Hofmann Lab for discussion and assistance.

## Ethical approval

The authors certify that this work followed ethical treatment of animals outlined in their IACUC protocol (AUP-2018-00236).

## Author’s Contributions

HAH and KJW conceived of the study. KJW designed and constructed the experimental apparatus and data collection procedures, collected behavioral data, and performed statistical analyses. HAH and KJW interpreted the results. KJW wrote the initial draft of the manuscript, and HAH provided feedback and comments during manuscript writing. All authors give final approval for publication.

