## Supplementary material for "Equal performance but distinct behaviors: *Astatotilapia burtoni* sex differences in a novel object recognition task and spatial maze": Electronic Supplementary Materials

Wallace & Hofmann 2020

**Electronic Supplemental Material**

---

**Table of Contents:**


**Page 2**- Supplemental Tables 1-3: *PCA loadings*

**Page 3**- Supplemental Figure 1: *Image of naturalistic pool community housing*

**Page 4**- Supplemental Figure 2: *Objects used in the NOR task (with dimensions)*

**Page 5**- Supplemental Figure 3: *Average space use in the NOR task over time (60 second segments) for females (left) and males (right)*

**Page 6**- Supplemental Figure 4: *In the fourth trial of the SPA task, female first choose the rewarded side significantly more than males.*

**Page 7**- Supplemental Figure 5: *Average space use in the SPA task over time (60 second segments) for females (left) and males (right)*

**Page 8**- Supplemental Figure 6: *Principal Component Analysis of the Novel Object Recognition Task.*

**Page 9**- Supplemental Figure 7: *Principal Component Analysis of the Spatial Task.*

**Page 10**- Supplemental Tables 4-5: *Selected Model and Model Averaged Importance of Terms Analysis for variables in the NOR task (4) and SPA task (5)*

**Supplemental Table 1.** NOR PCA Loadings

| Variable Name | PC1 | PC2 | PC3 |
|---|---|---|---|
| **NOR non-engagement** | 0.3337095 | 0.06198056 | 0.57247477 |
| **NOR familiar preference** | -0.2770051 | -0.51964999 | 0.17459412 |
| **NOR novel object approaches** | 0.2075964 | 0.34836716 | -0.24689154 |
| **NOR familiar object interaction bout duration** | 0.2233546 | -0.49244333 | -0.13605707 |
| **NOR familiar object approaches** | -0.2024670 | -0.39922392 | 0.05851764 |
| **NOR % interaction** | 0.4225524 | -0.34998333 | -0.35391929 |
| **NOR novel object interaction bout duration** | 0.3700800 | 0.14707399 | -0.26677032 |
| **NOR % association** | -0.5463550 | 0.23562313 | -0.08885502 |
| **NOR object approach latency** | 0.2528166 | 0.05696578 | 0.59546434 |

**Supplemental Table 2.** SPA PCA Loadings

| Variable Name | PC1 | PC2 | PC3 | PC4 |
|---|---|---|---|---|
| **SPA % interaction** | -0.24840614 | -0.42198824 | -0.45038831 | 0.10380760 |
| **SPA decision latency** | 0.05172298 | 0.44311654 | -0.47391578 | 0.02395896 |
| **SPA activity** | 0.35694550 | -0.12457042 | 0.10841213 | -0.28331353 |
| **SPA % association** | -0.02738082 | 0.42464177 | 0.53421119 | -0.29398232 |
| **SPA % decision areas** | -0.44633462 | -0.03270950 | 0.08980397 | -0.28379179 |
| **SPA reward preference** | -0.38066121 | 0.20596072 | 0.07374121 | 0.44389381 |
| **SPA thigmotaxis** | 0.46194293 | -0.07625763 | 0.02896399 | 0.27141066 |
| **SPA reward latency** | 0.10795448 | 0.20120429 | -0.47642436 | -0.59521126 |
| **SPA interaction latency** | 0.11987986 | 0.56792881 | -0.18133135 | 0.31600566 |
| **SPA reward preference following solution** | -0.47320731 | 0.13192415 | -0.03488214 | -0.12875765 |

**Supplemental Table 3.** Combined (NOR + SPA) PCA Loadings

| Variable Name | PC1 | PC2 | PC3 | PC4 |
|---|---|---|---|---|
| **NOR % association** | -0.19811801 | -0.332608625 | 0.06791586 | -0.04835448 |
| **NOR novel object approaches** | 0.03466594 | 0.311318169 | 0.33113801 | 0.12889048 |
| **SPA % interaction** | -0.27816596 | -0.144396164 | 0.28787844 | -0.22400223 |
| **SPA thigmotaxis** | 0.40003132 | -0.090449985 | 0.08687745 | -0.02021460 |
| **SPA activity** | 0.38507403 | -0.052510802 | 0.06006192 | 0.02240206 |
| **NOR familiar preference** | 0.09997752 | -0.409809490 | -0.35547884 | -0.04490277 |
| **NOR familiar object approaches** | 0.19411637 | -0.324627775 | -0.24040723 | -0.01403501 |
| **NOR % interaction** | 0.25718226 | 0.149740955 | 0.03684055 | 0.31047254 |
| **NOR familiar object interaction bout duration** | 0.22767278 | -0.118025764 | -0.29726884 | 0.11032418 |
| **SPA reward latency** | 0.06569466 | 0.053325024 | -0.06162624 | -0.54351331 |
| **NOR object approach latency** | -0.03202930 | 0.282169298 | -0.28406155 | -0.04378537 |
| **SPA decision latency** | 0.01074838 | 0.265542329 | -0.15591276 | -0.46900751 |
| **NOR non-engagement** | 0.06182646 | 0.282136034 | -0.10028802 | -0.14065924 |
| **SPA interaction latency** | 0.06245063 | 0.302303818 | -0.32924739 | -0.24342358 |
| **SPA % association** | 0.03961168 | 0.136575012 | -0.41720924 | 0.33359160 |
| **NOR novel object interaction bout duration** | 0.05779251 | 0.307084392 | 0.11452483 | 0.22325672 |
| **SPA % decision** | -0.37522297 | -0.029012821 | -0.14923352 | 0.12891652 |
| **SPA reward preference** | -0.30730059 | 0.114993333 | -0.19000345 | 0.20053252 |
| **SPA reward preference following solution** | -0.39356715 | 0.001145558 | -0.21215226 | 0.06994338 |

**Supplemental Figure 1.**

Image of naturalistic pool community housing

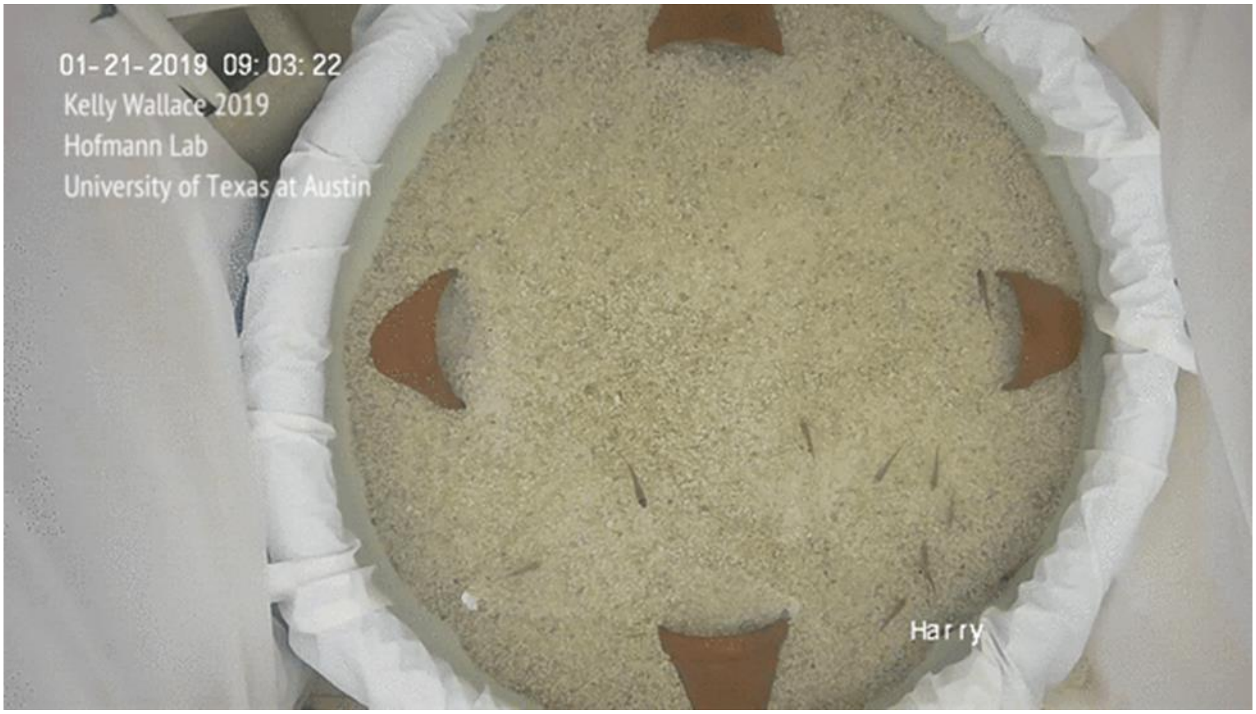

**Supplemental Figure 2.**

Objects used in the NOR task (with dimensions)

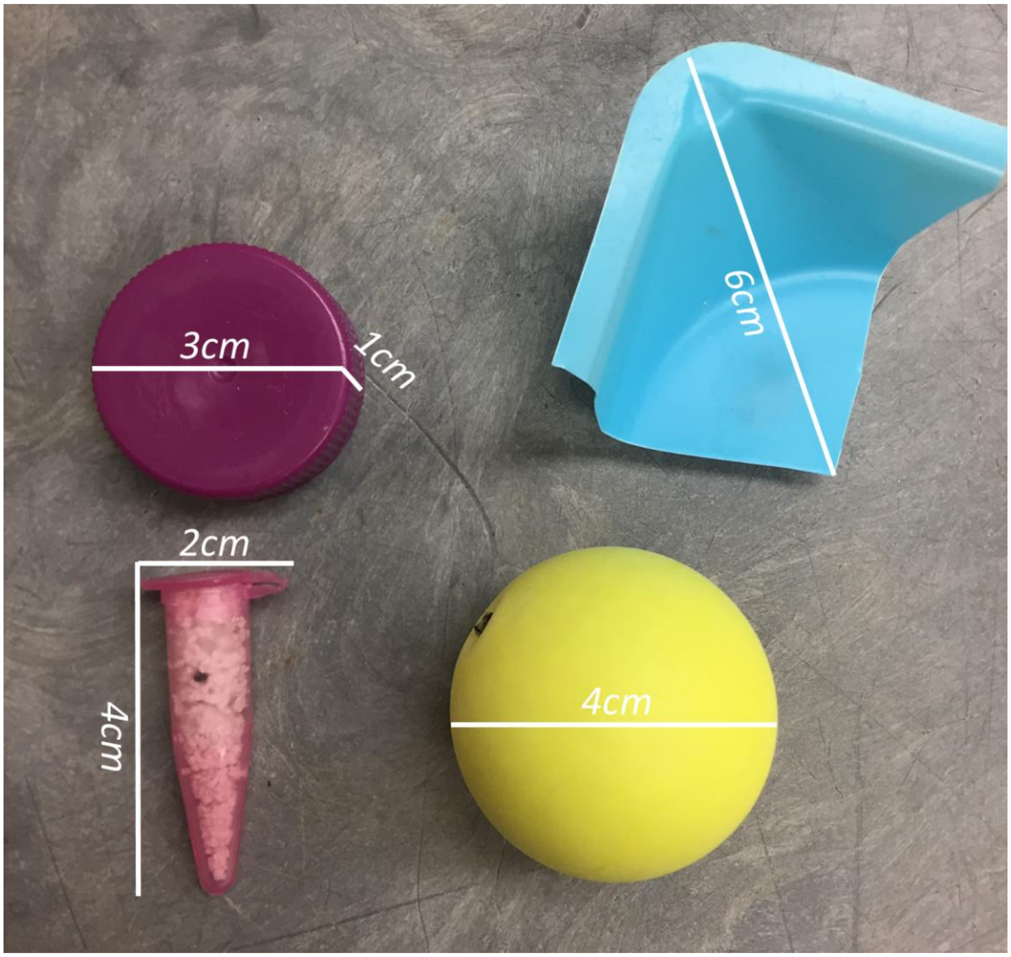

**Supplemental Figure 3.**

Average space use in the NOR task over time (60 second segments) for females (top) and males (bottom)

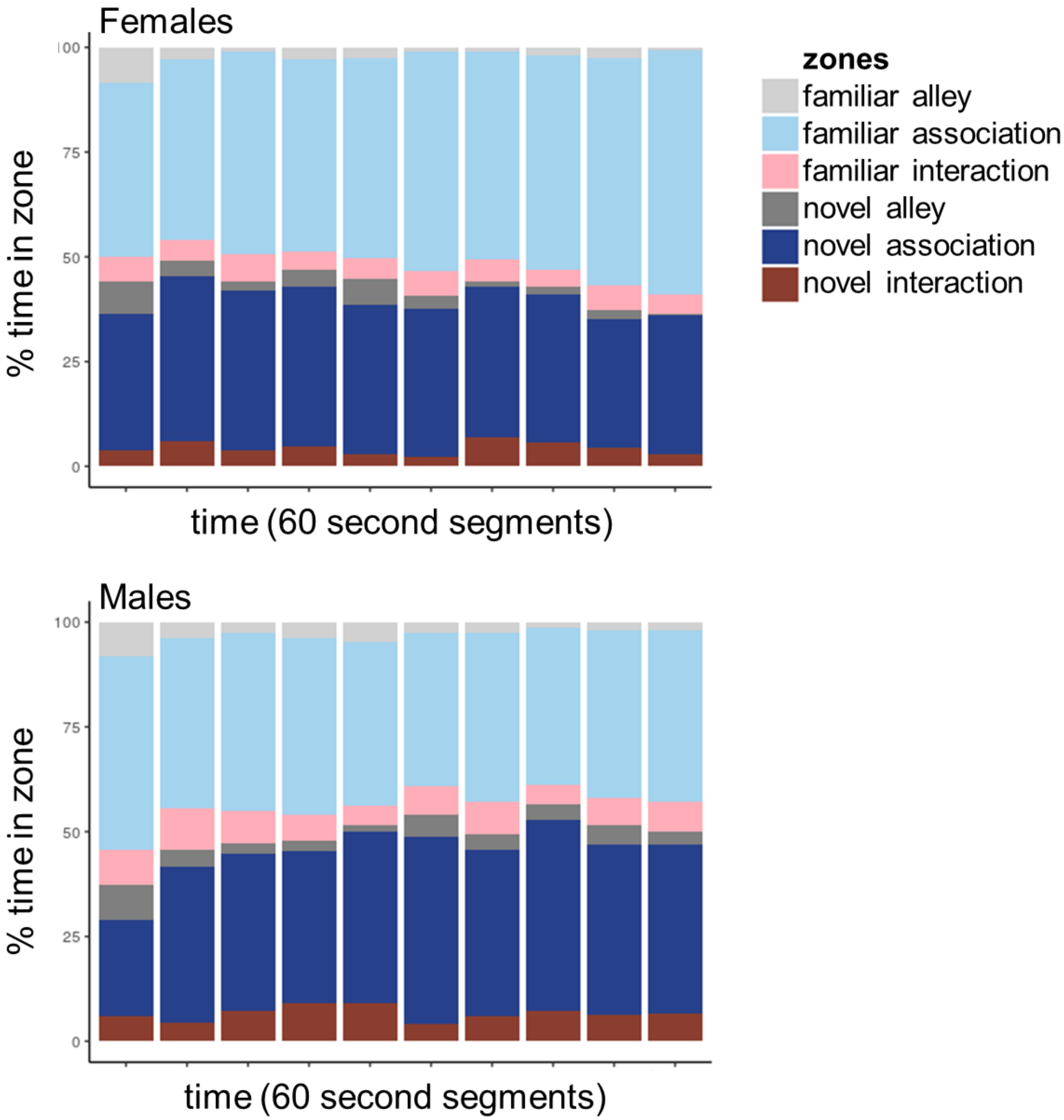

**Supplemental Figure 4.**

In the fourth trial of the SPA task, female first choose the rewarded side significantly more than males.

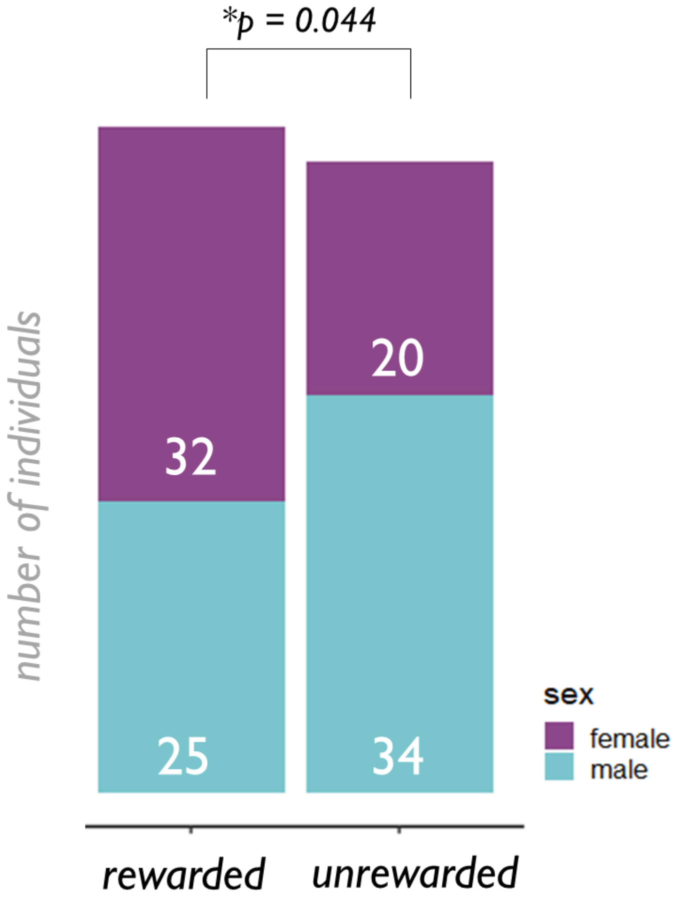

**Supplemental Figure 5.**

Average space use in the SPA task over time (60 second segments) for females (top) and males (bottom)

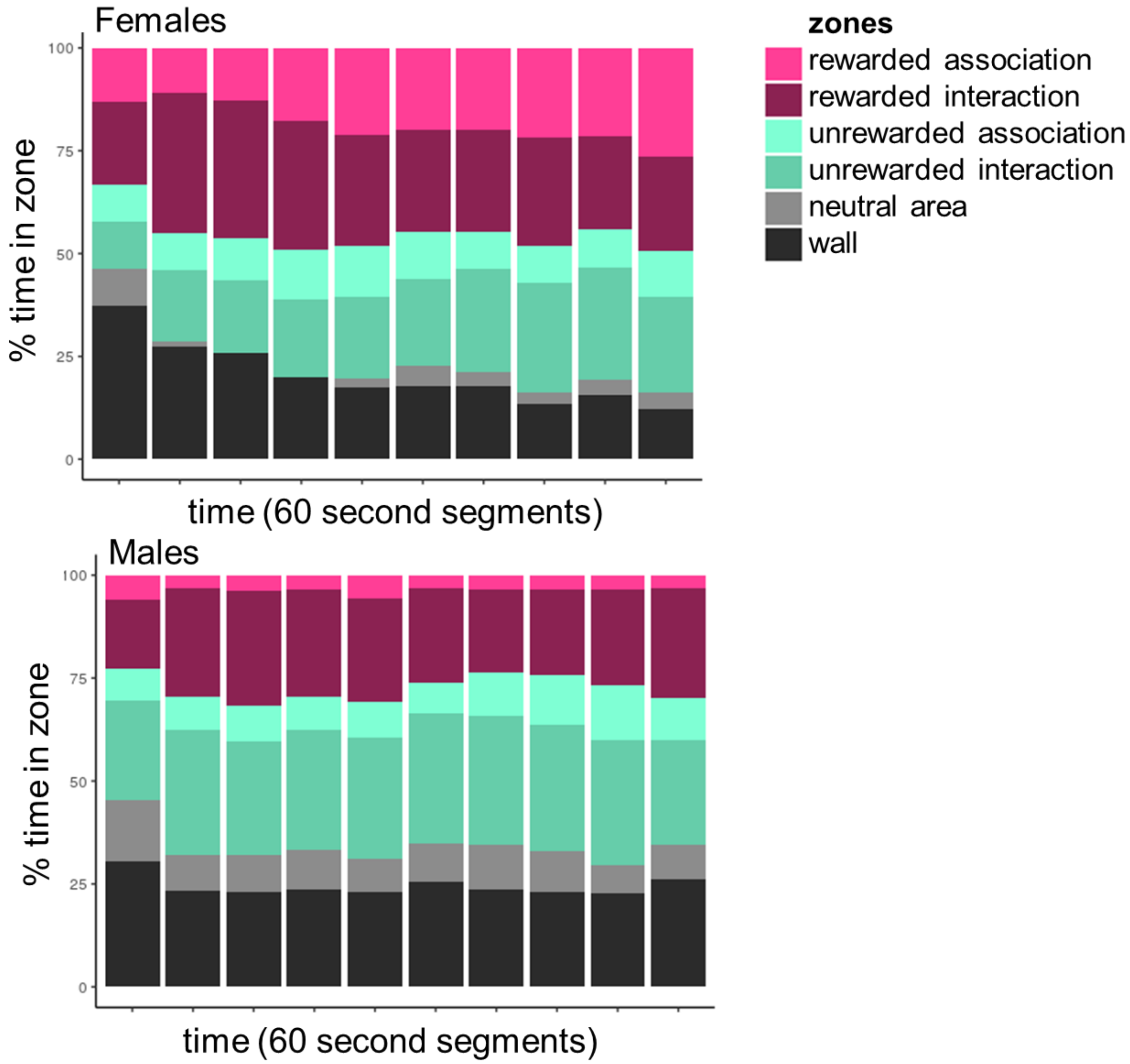

**Supplemental Figure 6.**

**Principal Component Analysis of the Novel Object Recognition Task.** A PCA of variables assessed in the NOR task identified three axes that explain >10% of the variance. Shown here are the proportion of variance for each PC axis (top) and a heatmap of the eigenvalues of each behavioral variable for each PC axis (bottom). The first PC axis (31%) can be described by variables of overall object association and interaction. The second PC axis (24%) can be described by familiar object preference. The third PC axis (19%) can be described by measures related to neophobia. A hierarchical clustering of variables by loading patterns (dendrogram) identified a highly supported cluster (AU > = 95%) of NOR familiar preference with NOR non-engagement (highlighted in grey box).

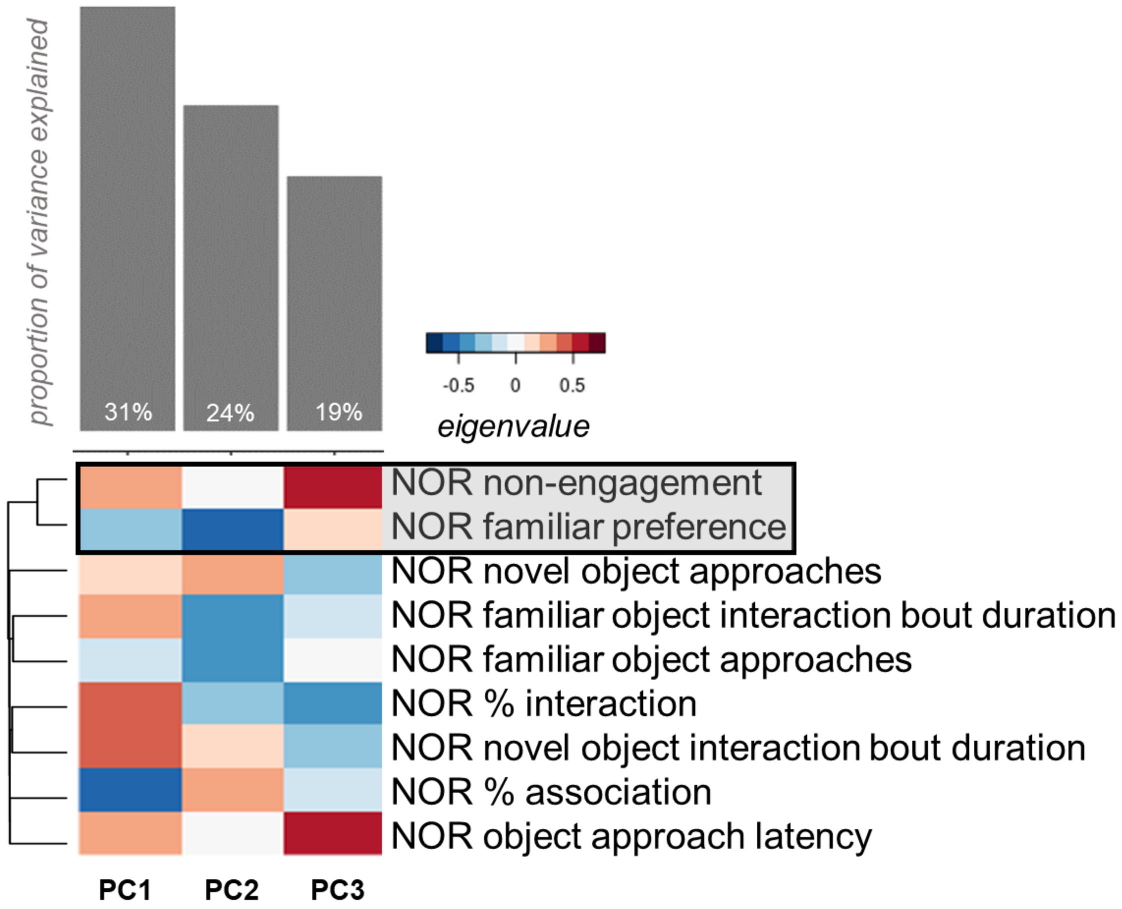

**Supplemental Figure 7.**

**Principal Component Analysis of the Spatial Task.** A PCA of variables assessed in the SPA task identifies four axes that explain >10% of the variance. Shown here are the proportion of variance for each PC axis (top) and a heatmap of the eigenvalues of each behavioral variable for each PC axis (bottom) (A). The first PC axis (36%) can be described by variables of preference and engagement. The second PC axis (20%) load most strongly by decision and interaction latencies. The third PC axis (18%) loads primarily with stimuli association. The fourth PC axis (11%) can be described by variables related to the reward stimulus (latency and preference). A hierarchical clustering of variables by loading patterns (dendrogram) identified a highly supported cluster (AU > = 95%) of NOR familiar preference with NOR non-engagement (outlined in black). PC2 significantly differs by sex (B).

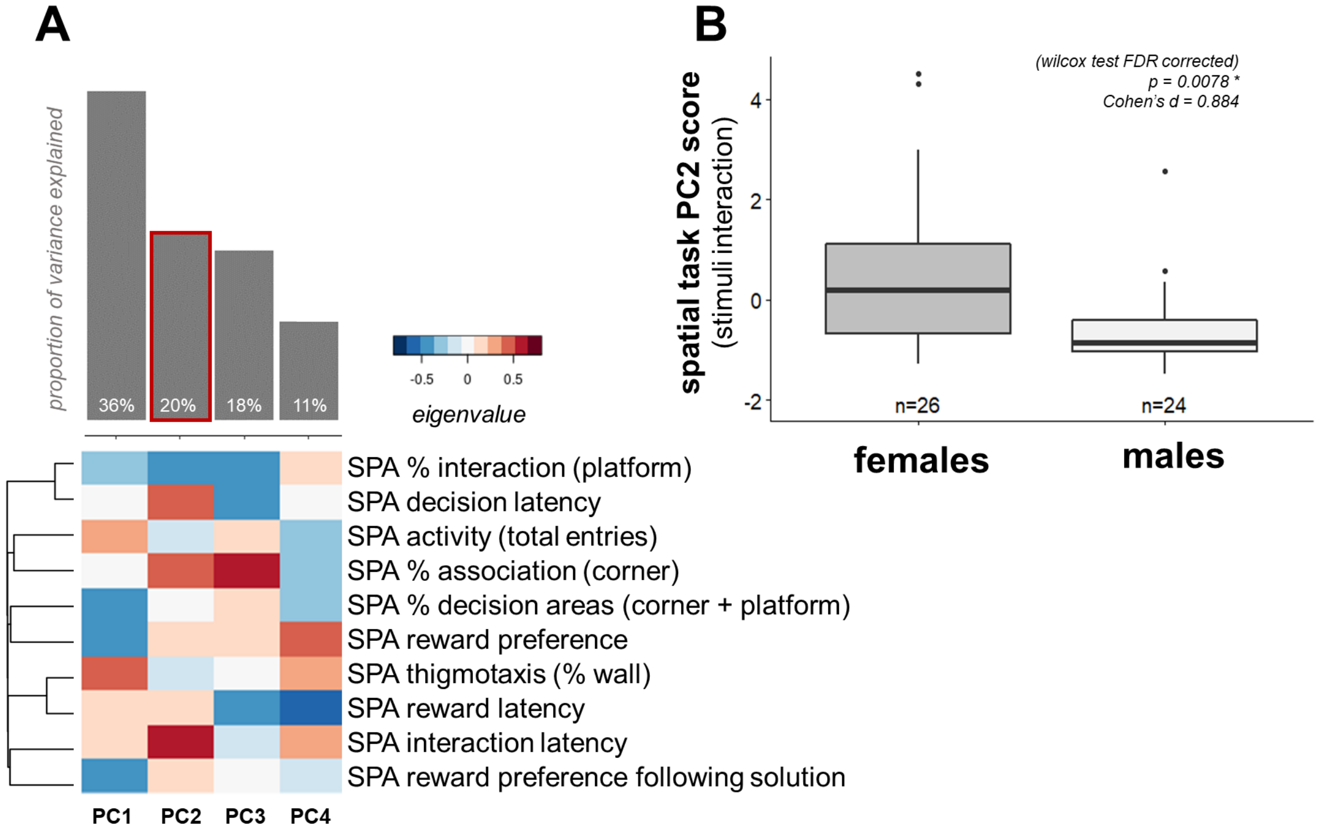

**Supplemental Table 4.** Selected Model & Model-Averaged Importance of Terms Analysis (NOR)

| Coefficients | Estimate | Std Error | z value | p |
|---|---|---|---|---|
| (Intercept) | -0.56 | 0.50 | -1.11 | 0.27 |
| NOR novel object interaction bout duration | 0.15 | 0.12 | 1.28 | 0.20 |
| **Selected Model Info & Fit** | | | | |
| Observations | 41 | | | |
| dependent variable | sex | | | |
| Type | generalized linear model | | | |
| Family | binomial | | | |
| Link Function | logit | | | |
| AICc | 59.307 | | | |
| McFadden's Pseudo-$R^2$ | 0.032 | | | |
| **Model Averaged Importance of Terms Analysis** | | | | |
| NOR novel object interaction bout duration | 0.5623127 | | | |
| NOR familiar preference | 0.4157714 | | | |
| NOR non-engagement | 0.3491010 | | | |
| NOR novel object approaches | 0.3341324 | | | |
| NOR % association | 0.3086410 | | | |
| NOR % interaction | 0.2246150 | | | |
| NOR familiar object approaches | 0.1569398 | | | |
| NOR object approach latency | 0.1561737 | | | |
| NOR familiar object interaction bout duration | 0.1438487 | | | |

**Supplemental Table 5.** Selected Model & Model-Averaged Importance of Terms Analysis (SPA)

| Coefficients | Estimate | Std Error | z value | p |
|---|---|---|---|---|
| (Intercept) | 3.12 | 1.32 | 2.37 | 0.02 |
| SPA reward preference | -3.02 | 1.43 | -2.11 | 0.04 |
| SPA % association | -4.45 | 2.05 | -2.18 | 0.03 |
| **Selected Model Info & Fit** | | | | |
| Observations | 41 | | | |
| dependent variable | sex | | | |
| Type | generalized linear model | | | |
| Family | binomial | | | |
| Link Function | logit | | | |
| AICc | 51.340 | | | |
| McFadden's Pseudo-$R^2$ | 0.213 | | | |
| **Model Averaged Importance of Terms Analysis** | | | | |
| SPA % association | 0.75444488 | | | |
| SPA reward preference | 0.43867812 | | | |
| SPA decision latency | 0.35158456 | | | |
| SPA % interaction | 0.33152292 | | | |
| SPA reward preference following solution | 0.32660784 | | | |
| SPA activity | 0.31213124 | | | |
| SPA % decision areas | 0.26853946 | | | |
| SPA interaction latency | 0.19728669 | | | |
| SPA reward latency | 0.19390881 | | | |
| SPA thigmotaxis | 0.06742759 | | | |
